# Simulations of flow over a bio-inspired undulated cylinder with dynamically morphing topography

**DOI:** 10.1101/2021.09.30.462620

**Authors:** Mikihisa Yuasa, Kathleen Lyons, Jennifer A. Franck

## Abstract

Inspired by the geometric properties of seal whiskers, the addition of spanwise undulations on a cylinder have been shown to lower the mean and oscillatory forces and modify the frequency of flow-induced vibration when compared to smooth cylinders. Previously, computational fluid dynamics (CFD) has been used to characterize the hydrodynamic response with respect to specific geometric features. However simulations are time intensive due to complex three-dimensional meshing and computation time, limiting the number of geometric perturbations explored. A method is proposed in which the specific geometric features of this complex topography can be modified during a simulation thereby decreasing the time per geometric modification and removing the need for manual meshing of the complex structure. The surface of the seal whisker inspired geometry is parameterized into seven defining parameters, each of which is directly controlled in order to morph the surface into any realization within the defined parameter space. Once validated, the algorithm is used to explore the undulation amplitudes in the chord and thickness directions, by varying each independently from 0 to 0.3 thicknesses in increments of 0.05 at a Reynolds number of 500. The force and frequency response are examined for this matrix of geometric parameters, yielding detailed force trends not previously investigated. The impact of the bio-inspired morphing algorithm will allow for further optimization and development of force-mitigating underwater devices and other engineering applications in need of vibration suppression, frequency tuning, or force reduction.

## 1. Introduction

Inspired by the unique geometry of seal whiskers, this paper computationally investigates the fluid flow over an undulated cylinder by actively morphing the geometry. Research has shown that seals are exceptionally skilled at detecting hydrodynamic disturbances, a skill which has been partially attributed to their whisker sensory system [1]. Unlike other mammals, seals have bumps along the span of the whiskers, forming a unique undulated cylindrical geometry [2]. Undulated cylinders inspired by the unique topography of seal whiskers are known to reduce drag and vortex-induced vibrations (VIV) by modifying the coherent wake structures typical of smooth cylinders [3]. These desirable features of undulated cylinders have made them intriguing design options for various engineering systems, such as hydrodynamic sensors, towing cables, support structures, and a variety of aerodynamic components. Furthermore, the ability to shift or morph the shape of undulations depending on variable flow conditions may be desirable for active flow control applications.

Due to their numerous engineering applications, the hydrodynamics of flow over whisker inspired geometries have been an active subject of research. Using digital photography measurements, Hanke et al. developed a model whisker described by seven unique geometric parameters, which contains two sets of undulations out of phase with one another, each periodic along the span of the whisker, as shown in Figure 1 [3]. Using a combination of computational and experimental methods, Hanke et al. confirmed that the von Karman vortex street, a regular pattern of alternating shed vortices, was replaced with a higher pressure and more symmetric wake, effectively eliminating VIV. In a subsequent experimental investigation, a direct comparison to an elliptical cylinder was performed using particle velocimetry coupled with dynamic mode decomposition of the flow field and indicated strong disruption of the energy redistribution process in the wake [4], which varied with respect to angle of attack [5, 6]. The response to flow disturbances has also been documented computationally [7] and experimentally [8].

**Figure 1:**
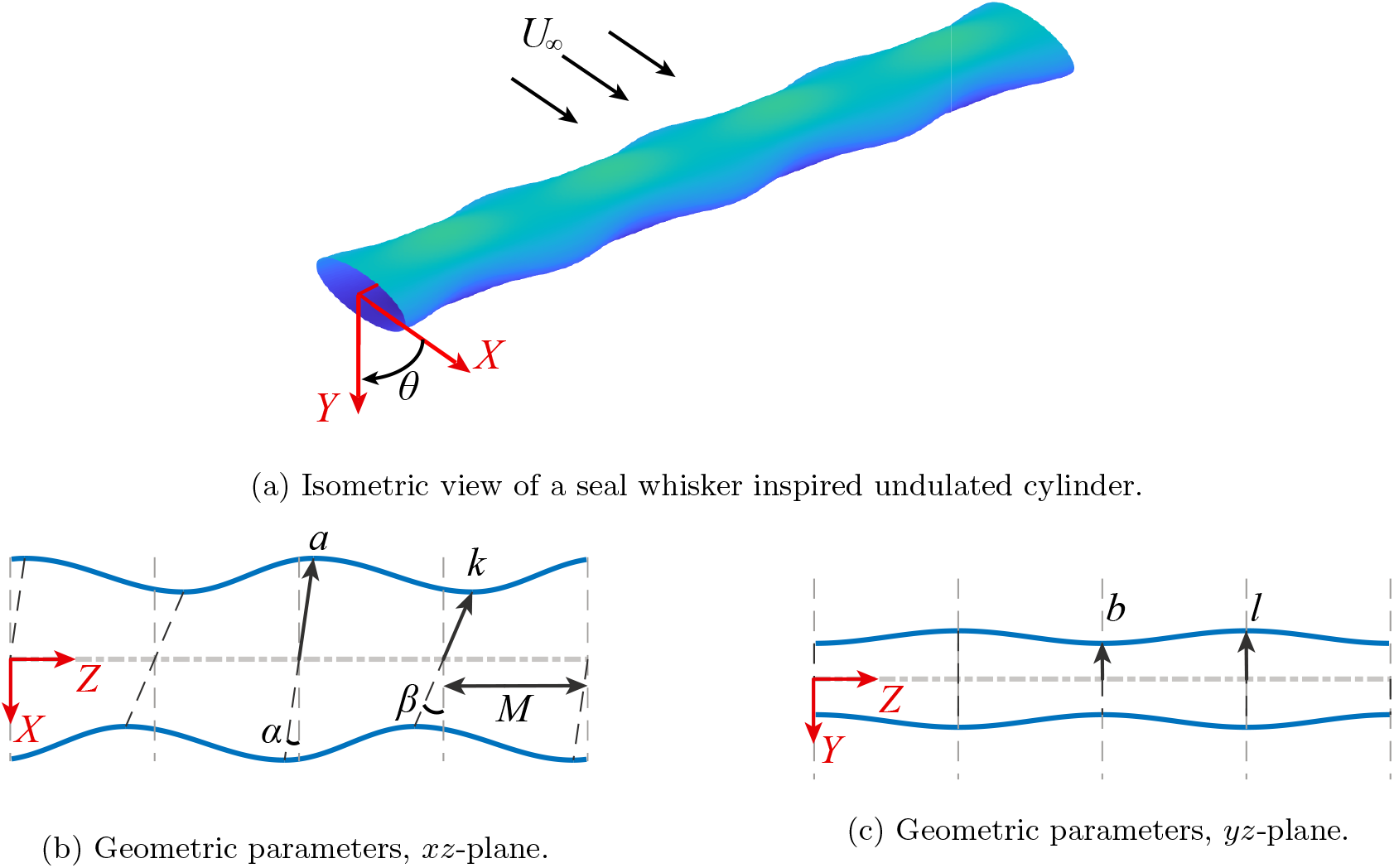
Isometric and cross-sectional views of undulated cylinder geometry, as defined by Hanke et al. [3]. Flow is in the positive *x*-direction.

The dual undulation geometric model described by Hanke et al. is a common method for defining the seal whisker geometry. The model is constructed based on average measurements of digital photography from 13 harbor seal whiskers. Building on work by Ginter et al [9] and Murphy [10], another independent investigation characterized mean geometric parameters from 27 whiskers, including both harbor and elephant seals [11]. It is shown that there is a wide statistical variability in whisker geometry, confirmed by morphological analyses that show geometric variation within and between species [2]. There have been several studies that explore the influence of geometric variation such as comparing two distinct wavelengths [12], removing one set of undulations [13], or shifting the phase between two sets of undulations [14]. Similar computational work has also been performed by Lam and Lin on wavy circular cylinders, in which the amplitude and wavelength are systematically varied [15]. Although not as complex as the seal whisker model, insight can be gained from their findings that show reductions in mean drag and lift oscillations with increasing amplitudes over a range of 0.02 to 0.3 diameters.

Recent work by Lyons et al. redefined the seven geometric parameters of the whisker model with respect to hydrodynamic relevance and explored a low and high perturbation from each nondimensional parameter from a baseline geometry [16]. A matrix of 16 geometric parameter combinations was created and a large-eddy simulation (LES) was used to assess the effect of each parameter on the overall flow response. The parameters that most significantly affect the flow response were found to be the aspect ratio, the wavelength, and the two undulation amplitudes.

Each of the previous computational investigations has been limited in the number of distinct models explored, making it difficult to assess the impacts of individual geometric parameters. Simulations are time intensive due to the three-dimensional geometry and inherent time-dependent and unsteady flow phenomena that must be resolved. Computational time is driven by the resolution and small time-step requirements, but there is also a large time requirement for the manual mesh generation of each new three-dimensional geometry proposed, including debugging for mesh errors and a long transient time to reach steady state before time-averaged statistics can be collected. Thus, computational studies consisting of numerous geometries must contend with compounding time constraints.

To alleviate the time and computational limitations, the current paper introduces a morphing whisker defined by the first complete topographic parameterization of the whisker surface. The analytic surface description is incorporated into a dynamic mesh morphing algorithm such that the geometric undulations are prescribed by the user throughout a single simulation, and the computational flow solver responds immediately to the changes in topography.

Using the newly developed method, the two orthogonal undulation amplitudes of the seal whisker model are modified systematically. Previous work has identified amplitude as an important parameter for drag reduction response [15, 16], and that the combination of two amplitudes in orthogonal directions is more beneficial than just one set of undulations [13, 17]. Recent research has also indicated that there may be a broad range of wavelengths and amplitudes at which the mean and fluctuating forces are reduced, perhaps beyond the morphological values found in seal whiskers [14]. To explore the amplitude parameter space in more detail, the undulated geometry is modified with the dynamic surface morphing technique. In the same manner as Lyons et al. [16], the two undulation amplitudes are recast into a nondimensional and hydrodynamically relevant chord length amplitude and thickness amplitude, allowing for one amplitude to be varied independent of another while holding the wavelength and phase-bias parameters constant. The results produce a matrix of seven chord length amplitudes by seven thickness amplitudes, for a total of 49 distinct undulated cylinder geometries. The flow response in terms of mean and fluctuating forces is discussed, linking it to the formation of unsteady flow structures and corresponding frequency spectra.

The algorithm and computational details are described in detail in Section 2. Section 3 compares the newly developed topography with previous results and documents the transient response of the flow to the mesh morphing process. Finally, the flow response to changes in two orthogonal undulation amplitudes is presented, highlighting changes to the forces, flow structures, and frequency spectra as a function of each undulation amplitude.

## 2. Methods

### 2.1. Overview of Mesh Morphing Strategy

Mesh deformation techniques are common in multidisciplinary optimization fields, especially those concerned with fluid flow around complex shapes and structures. Staten et al. detail the mesh morphing process for newly formed three-dimensional shapes encompassing a vast variation of geometries and outline the general process as follows: define the final mesh node positions, the curves connecting the nodes, and finally the surface mesh [18]. In a survey of techniques, Samreh et al. describe the various categories of mesh deformation, summarizing the advantages and disadvantages of each approach [19]. The method described in this paper falls under the basis vector approach, in which an analytical expression for the surface is developed and the changes from the initial to final state are prescribed by a sequence of vectors. The basis vector approach for dynamic meshing maintains the underlying structure of the mesh, which is beneficial for computational fluid dynamics (CFD) as it satisfies several requirements for fast simulation and convenient post-processing [20]. These benefits include the ability to consistently deform the target shape smoothly regardless of the complexity, utilize the minimum number of geometric variables to describe shape variations, maintain compatibility with existing geometries, and maintain stability of the solution. An example of the basis vector approach is implemented by Wei et al., who detail the motion of a structured mesh surface variation of square building corners, accomplished by moving mesh points in a single direction [21].

Taking a similar approach to Wei et al., the sequence of vectors describing the mesh motion of the undulated cylinder surface are confined to the radial direction, as illustrated in Figure 2. For simplicity, the original undeformed geometry is depicted as a smooth cylinder without any undulations in Figure 2a. However in practice, this initial geometry can be more complex and contain undulations defined by the prescribed analytical expressions. The underlying structured surface mesh contains *N_z_* points across the span and *N_θ_* points around the circumference. A vector, *r_o_*, defined from the *z*-axis, describes the location of each surface point. For the simple case of a smooth cylinder, the vector *r_o_* is a constant radius.

**Figure 2:**
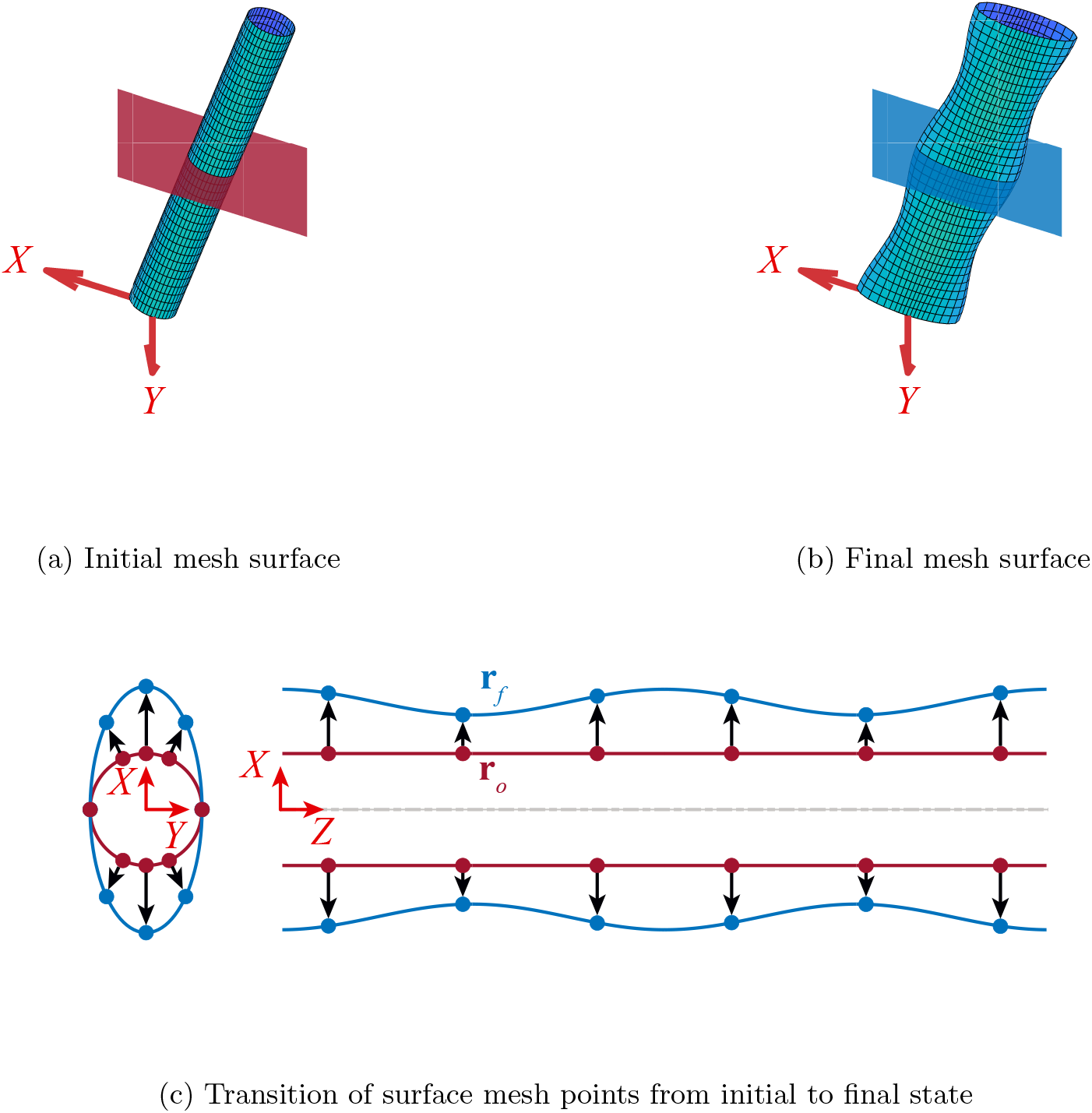
Overview of the mesh morphing from the initial to the final geometry based on the basis vector approach in which the surface points are confined to motion in the radial direction.

The final undulated cylinder in Figure 2b is defined to be spanwise periodic (along the *z*-axis) and each *xy*-plane maintains its *z*-coordinate throughout the morphing. Thus, points are constrained to move radially within the *xy*-plane throughout the deformation. The vector defining the final location of the deformed mesh point is given by *r_f_*, and the transformation vector is the difference between the final and initial shape. To ensure a smooth transition and stability of the fluid flow solver, the deformation is specified by multiple intermediate shapes by dividing the transformation vector into a sequence of vectors Δr_i_ [22], giving an expression for the final location as

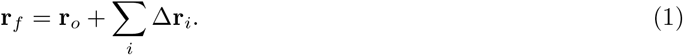

Moving the points radially ensures they maintain the same *θ* and *z* position before and after deformation. Thus, parameterization of the three-dimensional geometry is performed by examining the surface at each unique *θ*, as depicted in the *xz*-plane of Figure 2c. In this view, the surface geometry can be described as having a radius *r*, which is a function of *z* and the fixed parameter *θ*, or *r*(*z*; *θ*).

### 2.2. Parameterization of Undulated Cylinder Surface

Hanke et al. first characterized the whisker morphology by seven topographic parameters as shown in Figure 1: four semi-axes of the two inclined cross-sectional ellipses (*a*, *b*, *k*, *l*), two angles that define the ellipse inclination (*α*, *β*), and one distance parameter between the two ellipses (*M*) [3]. This definition contains inclined cross-sectional ellipses within the whisker geometry, only a skeletal structure of the geometry. Thus a complete analytical expression for the surface remains undefined. In order to morph the surface nodes into a new undulated geometry, the function *r*(*z*; *θ*) connecting the inclined ellipses must be defined with respect to the seven geometric parameters of the inclined ellipses.

The parameterized function for *r*(*z*; *θ*) is a two-part piecewise cosine function that intersects the geometry defining ellipses at points *R*_1_ and *R*_2_, as shown in Figure 3. As the geometry is rotated about the *z*-axis (thus changing the parameter *θ*), the radial and spanwise location of *R*_1_ and *R*_2_ move, but they remain the extrema of the function *r*(*z*; *θ*) by definition.

**Figure 3:**
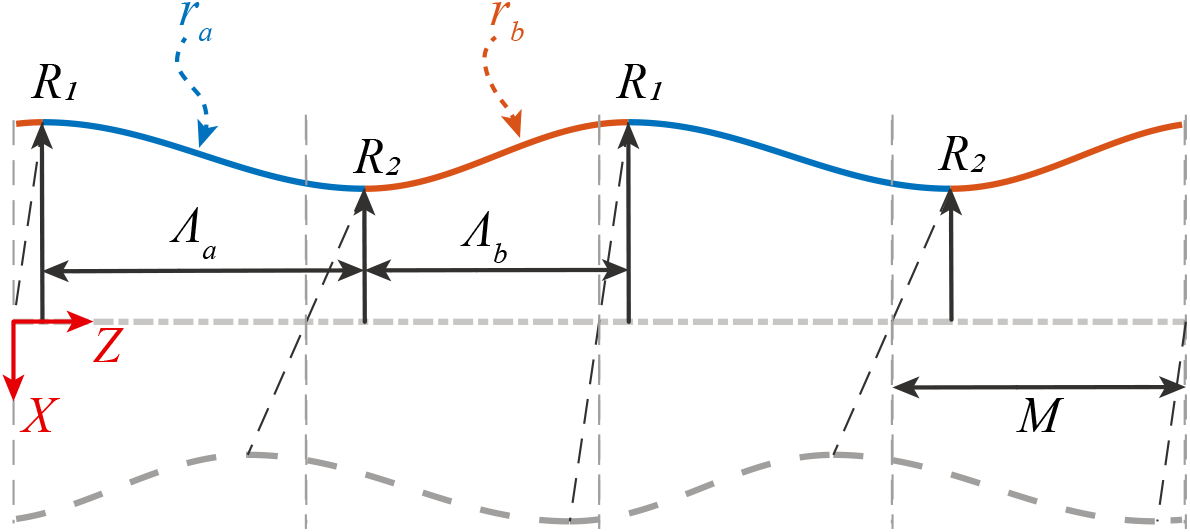
Schematic of the surface parameterization.

The radii *R*_1_ and *R*_2_ are calculated as

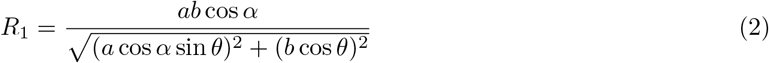

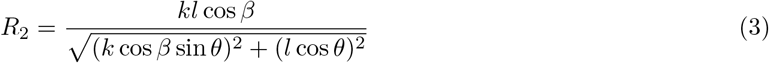

using the original ellipse parameters with a detailed derivation in Appendix Appendix A.1. The wavelengths of the cosine functions, Λ_α_ and Λ_b_, are similarly computed for each *θ* position as

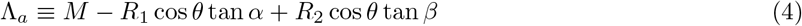

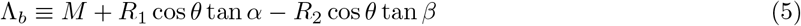

and are a function of *R*_1_ and *R*_2_, as well as the original ellipse parameters. Finally, the piecewise surface characterization function is expressed as

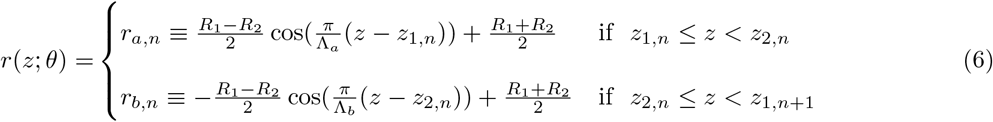

where

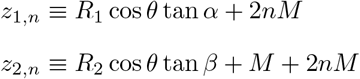

and 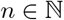.

Therefore, the surface vector r is expressed as r(*r*(*z*; *θ*), *θ*, *z*). The derivation above completes the analytical expression of the undulated whisker surface geometry based on the seven parameters that define the skeleton.

To morph between an initial and final geometry, a transformation vector is defined as r ≡ r_*f*_ – *r*_o_. Over a prescribed transformation time period, the surface experiences a smooth transition from the initial shape to the final shape using a time-dependent sigmoid function ς(*t*), scaled so that the transformation completes within the user-defined prescribed time period yielding

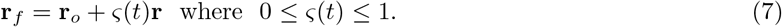

The smooth and gradual transition from the initial to the final shape described by Equation (7) is illustrated in Figure 4.

**Figure 4:**
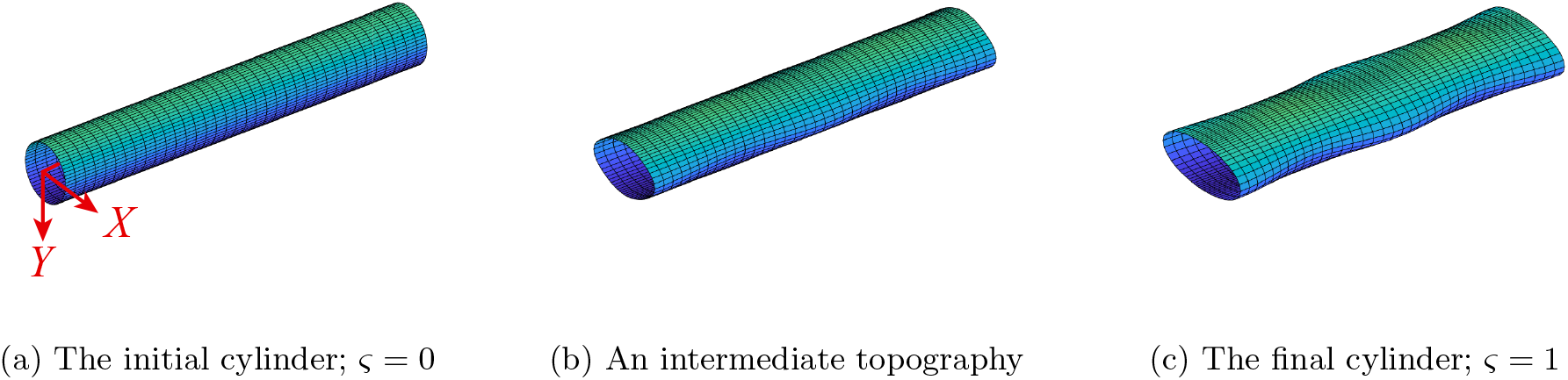
Transformation process using a time-dependent sigmoid function.

### 2.3. Computational Details and Implementation

The fluid flow solver utilizes the *OpenFOAM* finite volume libraries [23] to solve the incompressible Navier-Stokes equations,

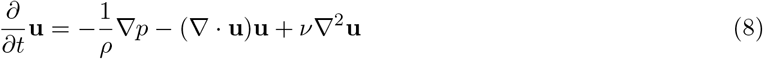

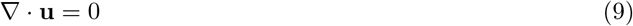

where u is the velocity vector, *ρ* is the density, *p* is the pressure, and *ν* is the kinematic viscosity. The fluid flow solver is coupled with the mesh motion [24] which solves the governing equation,

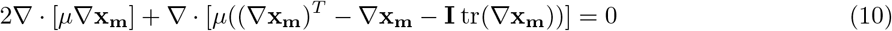

for the variable x_m_, or the displacement of each mesh node as a function of time. The variable *μ* represents a diffusivity in the mesh motion and is currently set to be inversely proportional to the distance from the mesh node to the cylinder surface. This setup ensures that the motion of the cells is spread out over the entire domain and not contained to the local deformation of the surface. Consequently, the mesh resolution near the surface, and thus within the boundary layer, does not change drastically as the cylinder evolves from one undulated topography to the next.

The dynamic mesh motion is driven through the boundary conditions at the surface of the cylinder, which are prescribed according to the equations presented in Section 2.2. As shown in Figure 5, the outer boundary is prescribed zero motion and remains stationary. During the morphing process, the mean thickness, *T*, of the cylinder remains constant, and is subsequently used as a reference length. The computational domain is circular in the *xy*-plane with a radius of 75T and extends two wavelengths in the spanwise direction, *L_z_*. Inlet flow boundary conditions are prescribed for *x* < 0, whereas outlet flow conditions are prescribed for *x* > 0, and periodic boundary conditions are applied across the span. A second-order finite volume method is utilized within *OpenFOAM* with a standard pressure-implicit split-operators (PISO) solver, and a second-order backward scheme is used to advance time.

**Figure 5:**
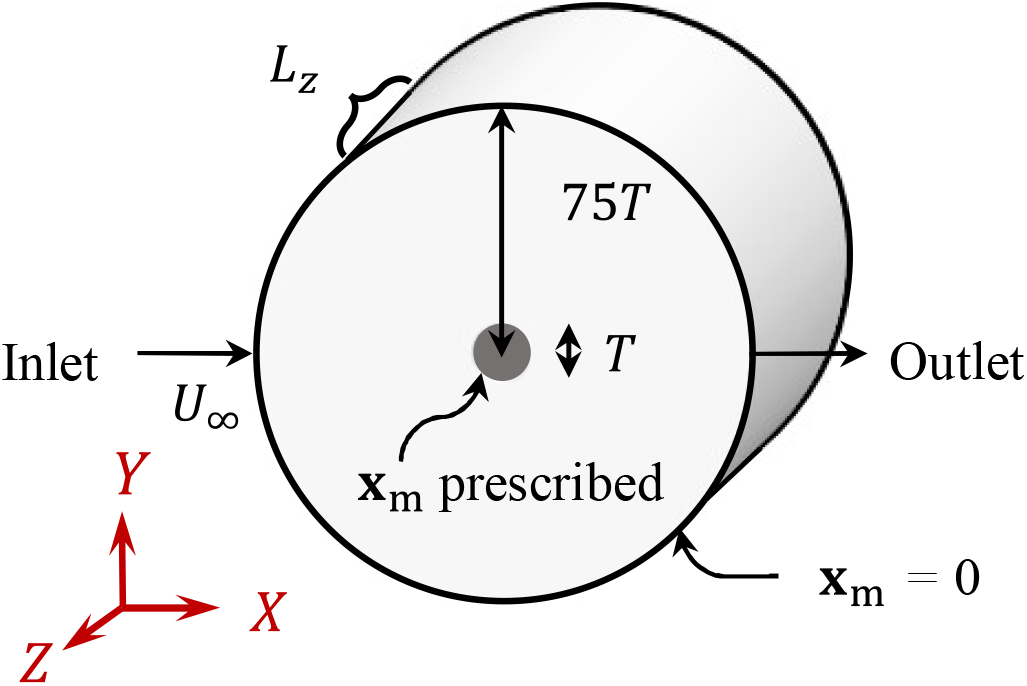
Schematic of the initial 3D computational domain with a smooth cylinder at center.

### 2.4. Simulation Setup

Fluid flow simulations are performed at an angle of attack of zero degrees such that the flow aligns with the *x*-axis, perpendicular to the cylinder thickness. As shown by Figure 1, the amplitude of the undulations are defined by *A_C_*, the amplitude in the chord or streamwise direction, and *A_T_*, the amplitude in the thickness or transverse direction. Both amplitudes are normalized by the mean thickness, *T*. Defining the amplitudes in this manner allows for each amplitude to be varied independently of one another while holding other hydrodynamically relevant parameters constant, such as the wavelength and aspect ratio. Changing the amplitudes *A_T_* and *A_C_* requires modification of six of the original geometric parameters, as *A_T_* = |*b* – *l*|/*T* and *A_C_* = |*α* cos *α* – *k* cos *β*|/*T*. A thorough explanation of the relationship between the amplitude-based geometric parameters of Figure 6 and the ellipse-based geometric parameters of Figure 1 can be found in Lyons et al. [16].

**Figure 6:**
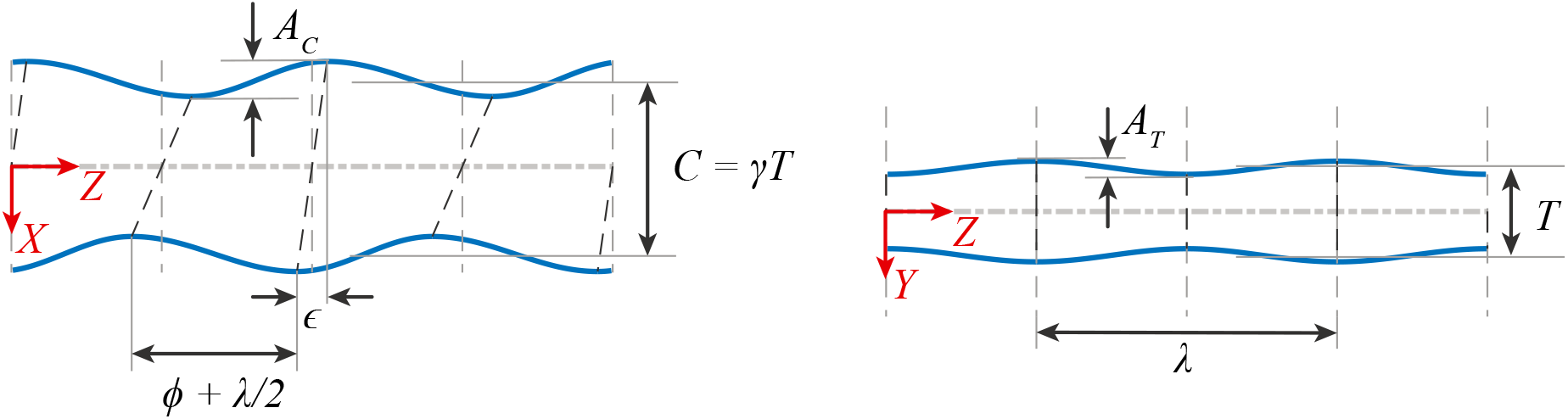
Geometric definitions of the amplitude in the chord length *A_C_*, amplitude in the thickness *A_T_*, the wavelength λ, aspect ratio *γ*, and the asymmetry parameters *ϵ* and *φ*.

To investigate the effect that changes in amplitude have on the flow response, each of the two nondimensional amplitudes is varied independently from 0 to 0.3 in increments of 0.05. The wavelength λ, nondimensionalized by thickness, is set to 3.43, and the aspect ratio *γ*, the ratio of the mean chord length to mean thickness, is set to 1.92. Both values are similar to dimensions measured in harbor seal whiskers [3, 9, 11]. The remaining two nondimensional quantities that introduce asymmetry into the model’s undulations as seen in whisker specimens are also held constant at *ϵ* = 0.342 and *φ* = 0.015, similar to values utilized in previous studies [3, 12, 13, 16, 17].

After each variation in geometry, the flow is allowed time to adjust to steady state conditions after which the flow response is averaged to compute the mean drag coefficient 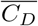, the root-mean-square (RMS) value of the lift oscillations *C_L,RMS_*, and the frequency spectra from the lift oscillations. The drag and lift coefficients *C_D_*, *C_L_* are computed as

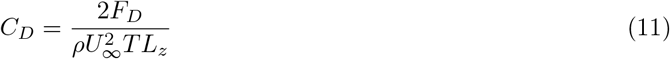

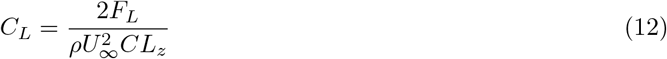

where *U*_∞_ is the freestream velocity, *L_z_* is the span length in the *z*-direction, and *ρ* is the fluid density. The drag and lift forces, *F_D_* and *F_L_*, are normalized by the averaged frontal and planform areas *TL_z_* and *CL_z_*, respectively.

The oscillating lift coefficient gives the frequency response *f*, which is nondimensionalized as

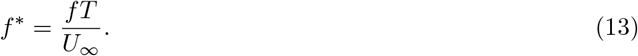

The most dominant peak in the frequency response is reported as the Strouhal number, *St*. Each new geometric perturbation is allotted 10 convective time units (CTU) for the transition, 150 CTU for the transient flow response once the shape has been formed and another 450 CTU for the steady state to collect the values of response variables. The Reynolds number of the simulations is nondimensionalized by the thickness, *T*, and is set to *Re_T_* = 500.

## 3. Results

### 3.1. Validation and Mesh Independence

The method described in Section 2.2 is, to the best of the authors’ knowledge, the first analytical parameterization of the entire seal whisker surface. The parameterization utilizes and maintains the definitions of the inclined cross-sectional ellipses first proposed by Hanke et al. and commonly used by researchers, and it also completely describes the periodic surface geometry between the ellipses. To assess the difference between the proposed analytical surface and surfaces created by other techniques, a comparison is performed with the topography in Lyons et al. [16]. In Lyons et al, the model is formed from two sets of inclined ellipses (for two wavelengths) and the whisker surface is created with the computer-aided-design (CAD) software *Solidworks* using a sequence of splines. Both models have the same nominal value of parameters *a*, *b*, *k*, *l*, *M*, *α* and *β*, although the interpolation between the inclined ellipses is performed differently.

To perform the comparison, the surface mesh from Lyons et al. is used as an initial configuration and the mesh morphing algorithm is subsequently applied. Thus, any radial motion of the mesh points will indicate differences between the new surface parameterization and the CAD generated surface. Figure 7a illustrates the distribution of the radial difference between each mesh point. The comparison shows that 96.6% of the mesh points are within 2% thickness difference with the original location. The maximum difference is 4.07% of the whisker thickness *T* and the mean and median difference are 0.59% and 0.41%, respectively. Figure 7b highlights the locations of larger radial coordinate differences across the two wavelengths. The largest differences can be observed near (*x*,*y*,*z*) = (−1, 0, 0.6) as well as (1, 0, 6.868) which is hidden due to the orientation in Figure 7b. The differences in Figure 7b also highlight that although the underlying ellipses form a periodic scaffold in the CAD implementation, this periodicity is not automatically applied in the *Solidworks* spline surface generation, leading to imperfect surface periodicity. The analytic function proposed in this paper alleviates this problem by mathematically describing the entire periodic surface.

**Figure 7:**
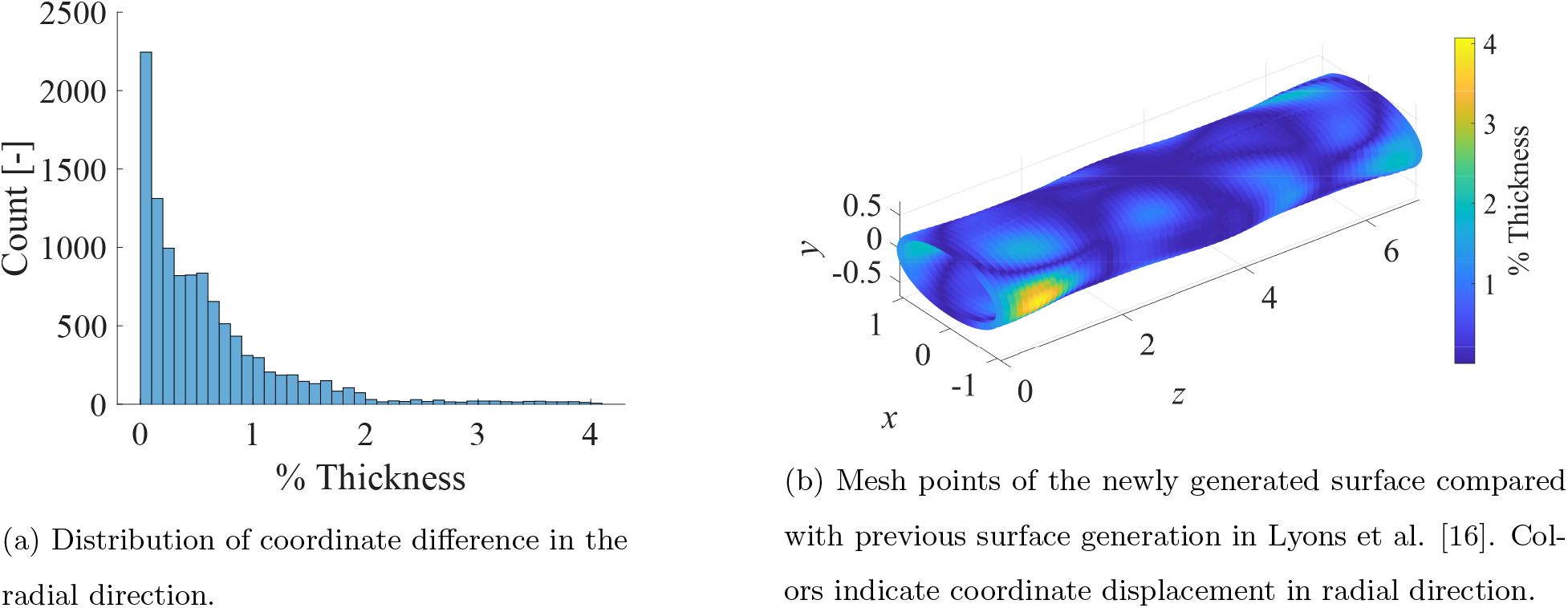
Illustration of (a) distribution and (b) locations of radial coordinate differences between a whisker surface generated by Lyons et al. [16] and by the newly proposed algorithm.

Mesh independence verification is performed by creating four smooth cylinder meshes of various resolutions. Each mesh is morphed into the same undulated whisker model geometry. The four different mesh resolutions are described in Table 1 in terms of the nodes in the radial, circumferential, and spanwise directions. As depicted in Figure 8, the points are clustered around the surface of the cylinder in the radial direction for resolution of the boundary layer (BL), clustered around the cylinder wake in the circumferential direction, and evenly spaced in the spanwise direction.

**Figure 8:**
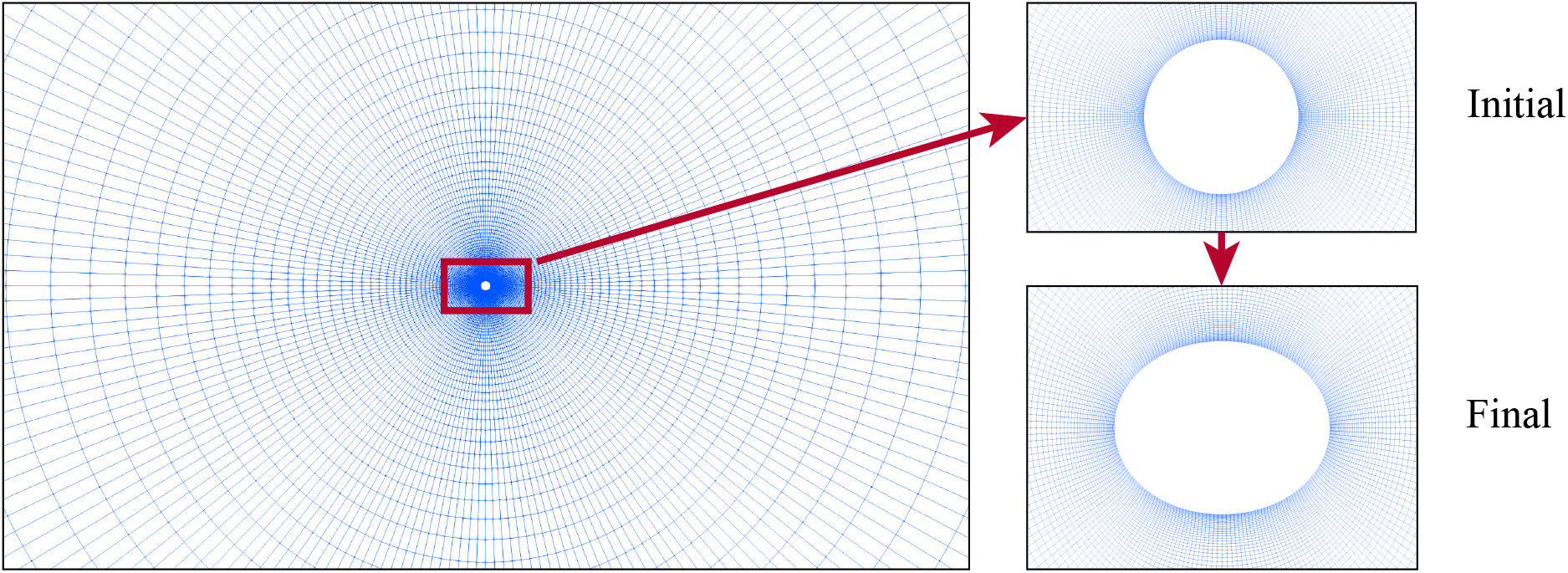
The *xy*-plane of the mesh with zoomed images of the mesh surrounding the initial smooth cylinder and the final mesh of the undulated cylinder.

**Table 1:** Resolution and mesh attributes of the underlying cylinder (initial configuration).

| Number of total mesh nodes $N$ | 2.57M | 1.92M | 1.39M | 0.718M |
| --- | --- | --- | --- | --- |
| $N_z$ | 110 | 100 | 90 | 68 |
| $N_\theta$ | 216 | 196 | 176 | 160 |
| $N_r$ | 110 | 100 | 90 | 68 |
| Mesh points in BL at $\theta = 90^\circ$ | 17 | 16 | 15 | 12 |
| $\Delta\theta_{\min}$ | 0.009561 | 0.01129 | 0.01331 | 0.01550 |
| $\Delta\theta_{\max}/\Delta\theta_{\min}$ | 3.137 | 2.678 | 2.254 | 1.935 |

The hydrodynamic properties (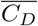, *C_L,rms_*) of the four different mesh resolutions in Table 1 are compared with one another and with those of previous investigations in Table 2. Each simulation is started from a fully developed circular cylinder flow then morphed over a time period of 10 CTUs. After waiting a transient period of 150 CTUs, the flow is averaged over the subsequent 450 CTUs to obtain the force response characteristics. As the resolution is increased, the mean drag slowly decreases to where there is only a 0.9% difference at the added expense of 0.65 million cells between the two most refined meshes, whereas the RMS lift force fluctuates between 0.0099 and 0.0122. The four mesh resolutions have comparable results with the previous studies at *Re_T_* = 390 – 500 despite the small changes in undulated geometry that will exist in the same manner that geometric differences between Lyons et al. and the current simulations are described in Section 1. Based on these results, and to best conserve the computational time, the mesh resolution of 1.92M is selected to carry out the current simulations.

**Table 2:** Mesh configurations of the undulated cylinder model compared with previous literature. Reynolds number from previous literature converted to be based on mean thickness, *T*, for consistency.

| | $N$ | $Re_T$ | $\overline{C_D}$ | $C_{L,RMS}$ |
| --- | --- | --- | --- | --- |
| 0.718M | 0.718M | 500 | 0.741 | 0.0157 |
| 1.39M | 1.39M | 500 | 0.714 | 0.0129 |
| 1.92M | 1.92M | 500 | 0.708 | 0.0099 |
| 2.57M | 2.57M | 500 | 0.701 | 0.0122 |
| Lyons et al. [16] | 1.92M | 500 | 0.717 | 0.0111 |
| Lyons et al. [16] | 0.718M | 500 | 0.715 | 0.0163 |
| Witte et al. [12] | 8M | 390 | 0.769 | 0.0173 |
| Hans et al. [13] | N/A | 390 | N/A | 0.011 |
| Kim and Yoon [6] | 3.5M | 390 | 0.741 | 0.0167 |

### 3.2. Transient response

Dynamic mesh morphing allows for a decrease in both user intervention and simulation time. Figure 9 compares the mesh morphing workflow (red) and a standard manual meshing workflow (blue), with an estimate on the manual hours and CPU hours based on the experience of the authors. In the standard workflow, a new mesh must be created by the user for each new geometry desired, even if the modification is minor. This method requires repeating the manual mesh debugging and transient flow development for each iteration. In contrast, dynamic mesh morphing can be implemented during an active simulation to transform the model from one topography into the next without user intervention. This method also decreases simulation time due to the faster transient regimes. Transitioning from one geometry with fully-developed flow to another geometry with fully developed flow is computationally faster than transitioning from an initial condition of uniform flow at all cells.

**Figure 9:**
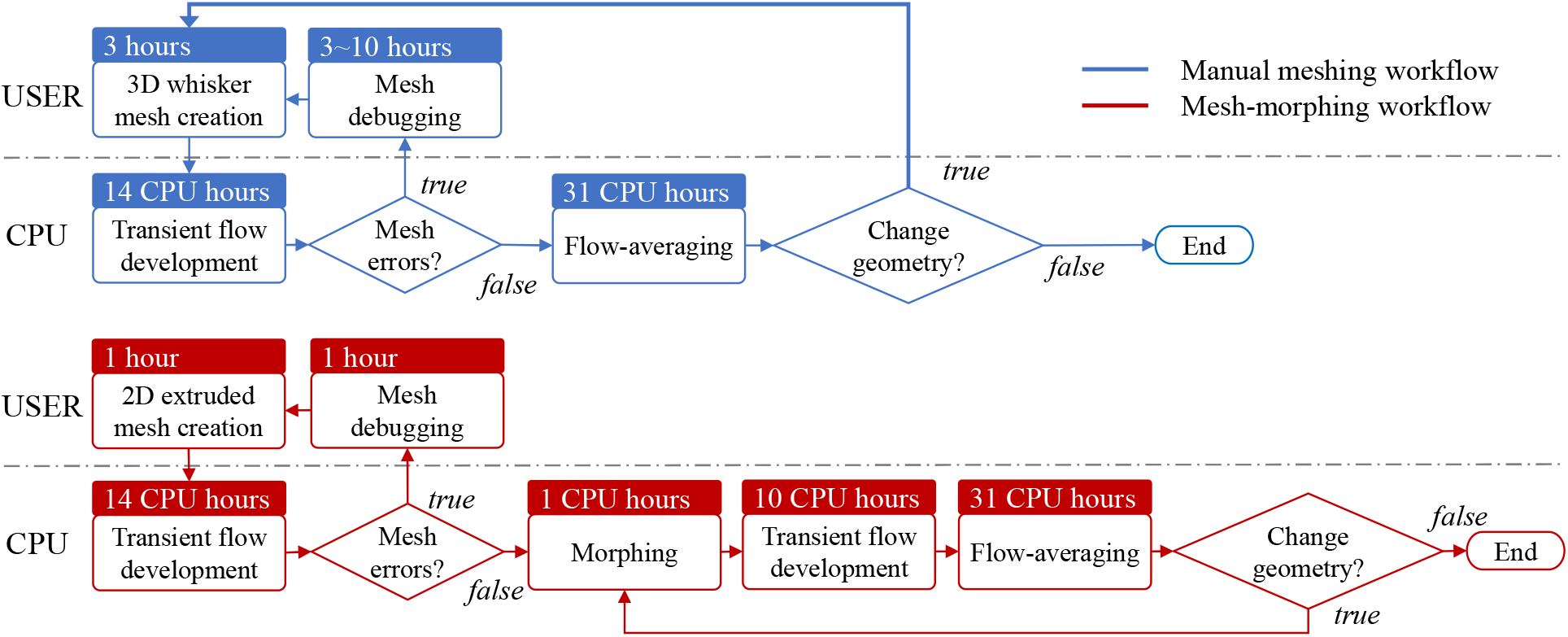
Comparison between a traditional mesh-solve-repeat process (top, in blue) and the proposed mesh-solve-morph process (bottom, in red).

An additional benefit to the mesh morphing workflow is that the initial mesh can be much simpler, in this case a two-dimensional (extruded) circular cylinder. A simpler mesh allows for a reduction in time for mesh creation and debugging. Due to two-dimensional and radially symmetric features, a circular cylinder mesh is easy to refine and validate. Subsequent morphed meshes are guaranteed to be structured due to the implementation of the morphing strategy.

To demonstrate the flow response to the dynamic motion of the mesh, a geometric transformation is performed with four undulated cylinder models where *A_T_* = 0 and *A_C_* increases through the values *A_C_* = 0.0, 0.1, 0.2, and 0.3, as shown in Figure 10. The hydrodynamic forces are recorded, showing the regions of active mesh morphing in red. From the response seen in both the drag and lift of Figure 10, the flow quickly adapts to new geometries and achieves a new steady state. The flow is allowed to develop around the new geometry for 150 convective time units (shown in green). The steady state regime (shown in blue) is where the time-averaged force statistics are computed.

**Figure 10:**
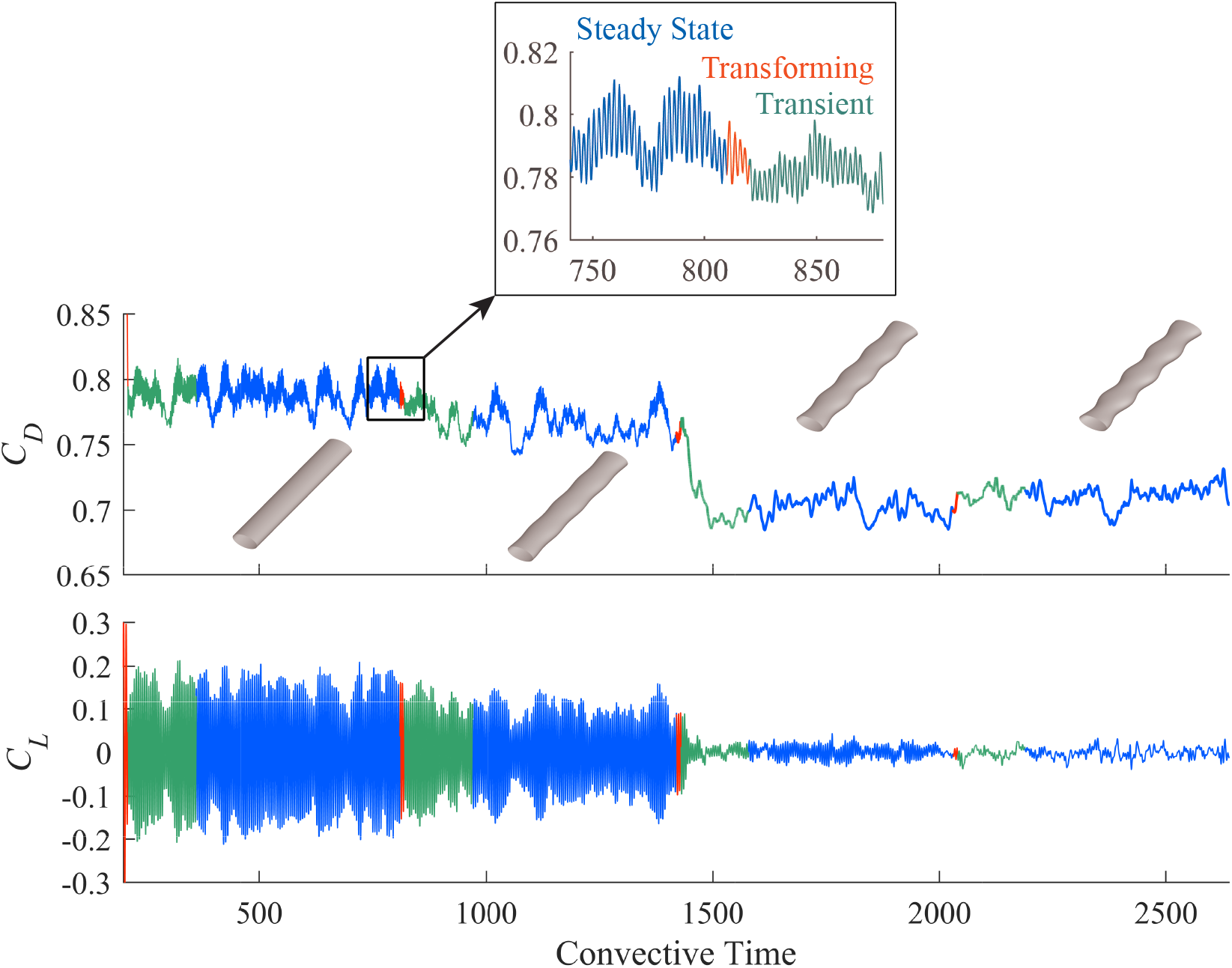
Results from a single simulation including mesh-morphing transitions between four models are shown. Models have a fixed value of *A_T_* = 0 with *A_C_* = 0, 0.1, 0.2, and 0.3. Plots of drag coefficient (top) and lift coefficient (bottom) show smooth mesh morphing transitions between each of the four geometric models.

In a rough estimate of the time savings for this four geometry simulation, using the values in Figure 9, the manual meshing and debugging process is reduced from 24 hours to 2 hours. This savings would only grow if more modifications are added, as there is no additional time cost for remeshing. Both methods take an equal amount of CPU time for four geometries, however the mesh-morphing becomes slightly more efficient as the number of geometries increase. This savings could be improved with less transient flow development which may be suitable for various applications.

### 3.3. Influence of Amplitude on Forces

To explore the effects of geometric modifications to a whisker inspired undulated cylinder, the two amplitude parameters are varied independently between 0 and 0.3 while holding the remaining geometric parameters constant, forming a matrix of 49 undulated cylinder geometries. The dynamic mesh morphing algorithm is utilized to reduce the 49 geometries to eight simulations, as shown in Figure 11, with each simulation containing six or seven unique topographies.

**Figure 11:**
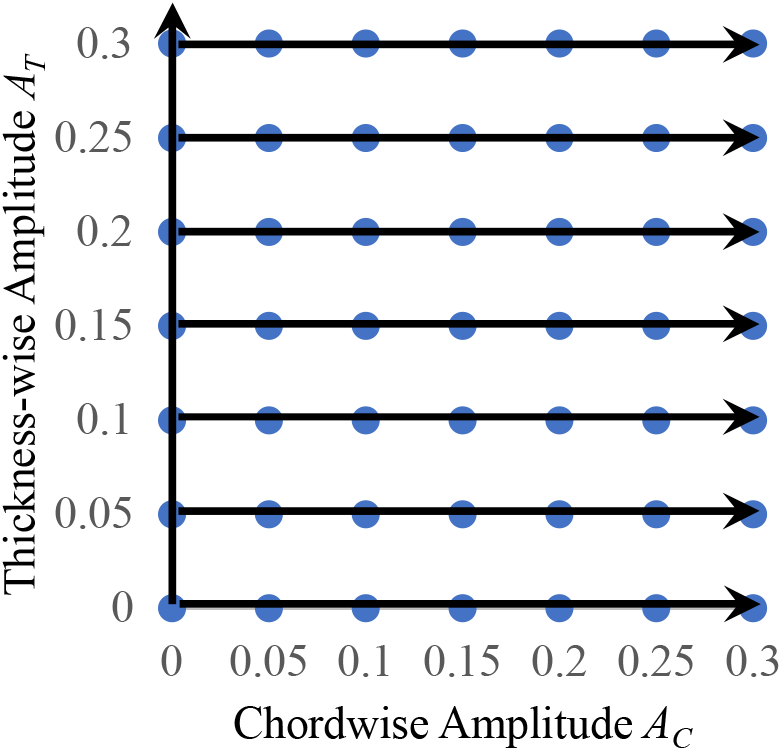
Parameter space of *A_C_* and *A_T_* variation. Each solid line denotes one simulation with multi-step transformations using the mesh morphing algorithm.

The hydrodynamic response for each of the 49 whisker geometries is quantified by 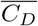 and *C_L,RMS_* with results shown in Figure 12. Both 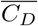 and *C_L,RMS_* show similar trends with respect to *A_C_* and *A_T_*, with maximum values in the lower left corner where *A_T_* and *A_C_* are zero, indicating the smooth elliptical cylinder topography. When *A_T_* = 0, the drag and oscillating lift coefficients decrease slightly as *A_C_* increases from 0 to 0.1. However, a larger drop is seen during the transition from an *A_C_* of 0.1 to 0.2. Similarly when *A_C_* = 0, 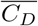 and *C_L,RMS_* decrease as *A_T_* increases from 0 to 0.3. However, the forces are reduced more quickly with the introduction of *A_T_* than *A_C_*. The response suggests the potential for a limiting threshold value for *A_C_* and *A_T_* if oscillating lift and drag reduction is desirable.

**Figure 12:**
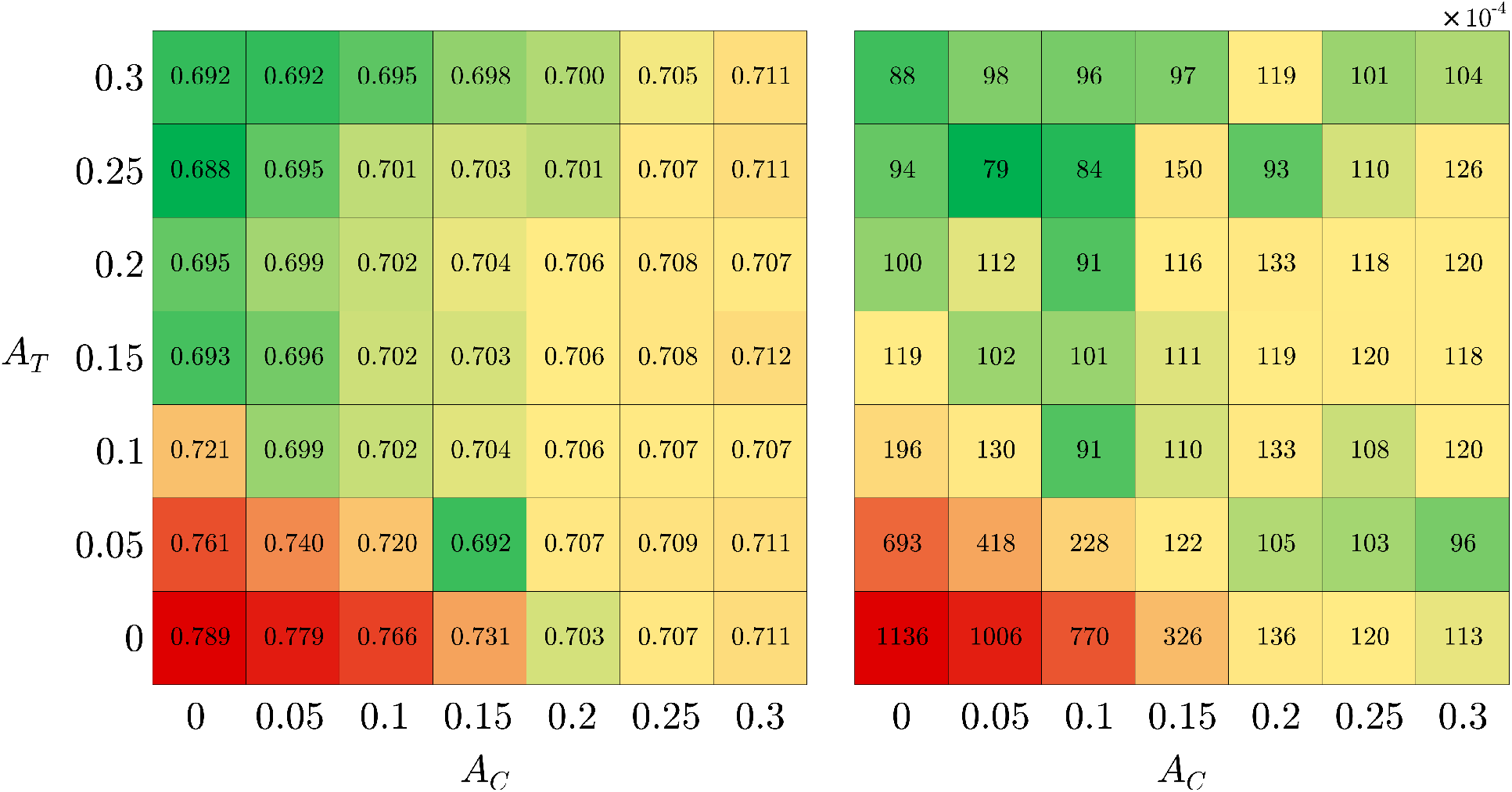
Flow response in terms of 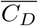 (left) and *C_L,RMS_* (right) for the 49 geometric variations of amplitudes.

At low to moderate *A_T_* values, *A_T_* = 0.05 and 0.1, *C_L,RMS_* continues the previous pattern of reduction with increasing *A_C_*. The drag initially decreases as *A_C_* increases with local minima at (*A_C_*, *A_T_*) = (0.05, 0.1) and (0.15, 0.05) then begins to increase again at larger *A_C_* values but remains lower than the absolute maximum at (0, 0). When *A_T_* ≥ 0.15, 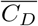 generally monotonically increases with larger *A_C_* values; however, the *C_L,RMS_* response is more complex. For *A_T_* ≥ 0.1, the local minima occur when *A_C_* = 0.05 — 0.1. Finally when *A_T_* = 0.3, *C_L,RMS_* is consistently low from *A_C_* = 0 – 0.15, giving a wider region of low lift oscillations in the top left corner, and a global minimum at (*A_C_*, *A_T_*) = (0.05, 0.25).

Although it is clear that small changes in *A_T_* and *A_C_* affect the flow, the dominant effects are examined by looking at flow features from the extreme values, (*A_C_*, *A_T_*) = (0, 0.3), (0.3,0) and (0.3, 0.3). Given the changes in spanwise undulations across the models, notable differences can be seen in the spanwise velocity component, *W*. Figure 13a displays the spanwise velocity of the these three cases within two *yz*-planes, one near the leading edge and another just before the trailing edge of the geometry. In the (0,0.3) case, there are large undulations in thickness and thus the freestream flow is quickly redirected into the positive and negative *z*-direction to overcome the change in thickness, displaying strong (10% of freestream) spanwise velocity immediately at the leading edge. Flow also continues to accelerate in the *x*-direction as shown by the streamlines in Figure 13b. The early onset of the spanwise velocity component gives way to alternate bands of *y*-vorticity forming at the leading edge and continuing to grow along the surface before being transported into the wake as seen in Figure 13c. Interestingly, the spanwise velocity is lower at the trailing edge than the leading edge. Of the three cases demonstrated, this has the lowest mean drag and the lowest RMS lift.

**Figure 13:**
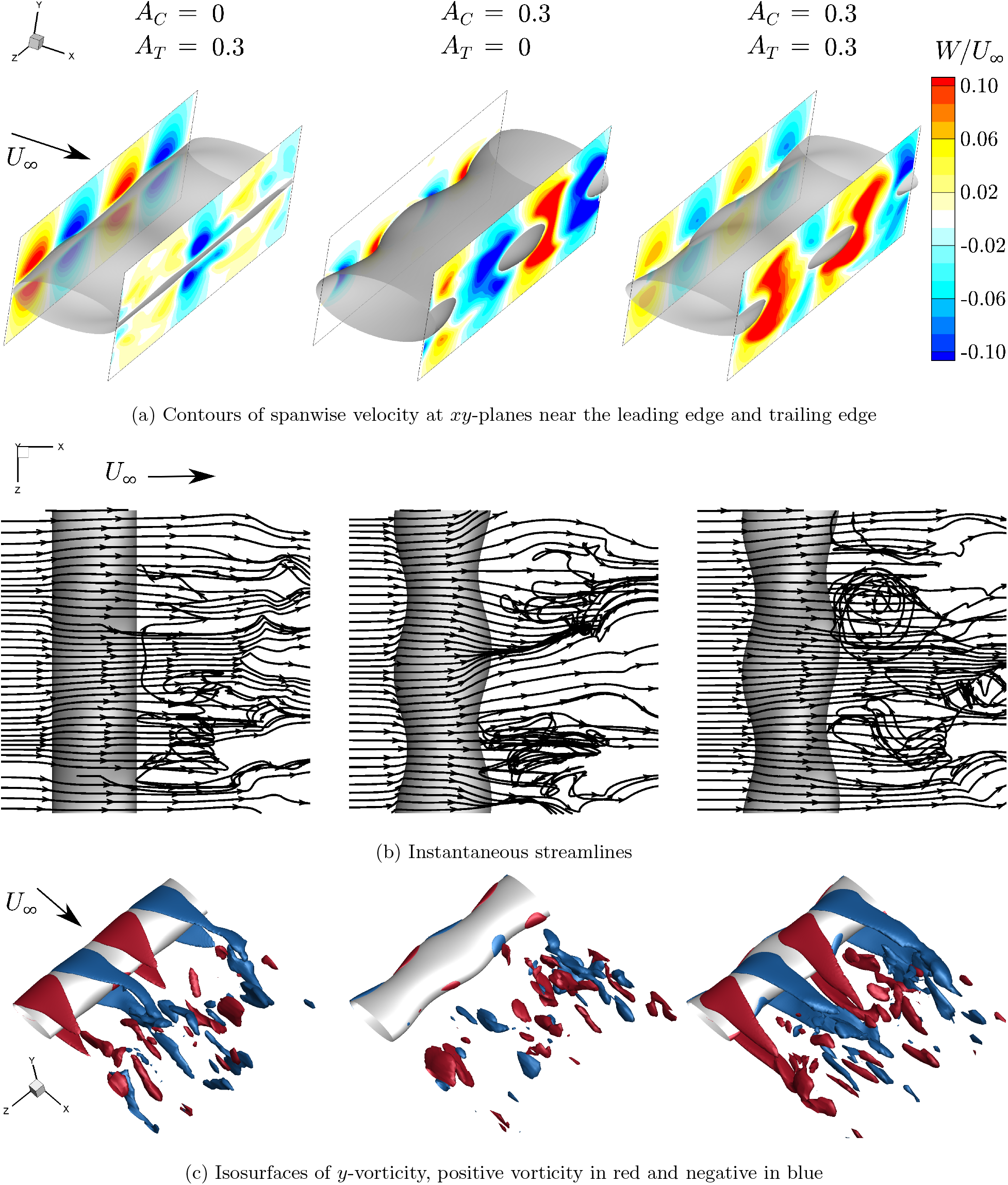
Comparison among simulated cases with extreme amplitudes, (*A_C_*, *A_T_*) = (0, 0.3), (0.3, 0), and (0.3, 0.3), shows the variation in induced spanwise flow and subsequent *y*-vorticity development.

In the second column is (*A_C_*, *A_T_*) = (0.3, 0), corresponding to a high amplitude in chord and no undulations in thickness. In contrast to the first model, the (0.3, 0) geometry produces a minimal amount of spanwise velocity at its leading edge. Spanwise flow develops as it travels over the surface, and has increased substantially by the trailing edge, revealing large regions of alternating positive and negative flow with *W* > 0.1 U_∞_. This later development of *z*-velocity is a consequence of the spanwise pressure gradient induced by the change in local chord length. As flow moves over the shorter chord length cross-sections, a larger adverse pressure gradient is created than for the more streamlined (long chord length) cross-sections. Thus, the flow shifts towards the low pressure region near the long chord length trailing edge as can be visualized by the streamlines in Figure 13b which seem to accumulate around the locations of highest amplitude at the trailing edge. In contrast to the high thickness amplitude model, this process develops very little *y*-vorticity along the surface.

In the third column of Figure 13, (*A_C_*, *A_T_*) = (0.3,0.3) demonstrates a super-positioning of two large amplitudes in both chord and thickness. The initial spanwise velocity is muted due to the presence of the chord length amplitude, and the tendency of the streamlines to gather at the highest chord amplitude is diminished with the introduction of the thickness amplitude. Nevertheless, the (0.3, 0.3) model develops *y*-vorticity early and then appears to sustain the layers further downstream as the spanwise flow feeds into them at the trailing edge. This most extreme combination of amplitudes still has a moderate drag reduction compared to other models tested matching that of the (0.3, 0) model, and a low to moderate RMS lift reduction, performing better than (0.3, 0).

### 3.4. Influence of Amplitude on Frequency Response

Introduction of three-dimensional vorticity as shown in Figure 13c can help prevent a dominant twodimensional spanwise shear layer roll-up, a phenomena previously noted by others [15, 14]. In addition to decreasing the amplitude of the lift response, the change in vortical shedding patterns directly affect the frequency distribution of the lift signal. An analysis of the lift signal spectra provides insight into the flow characteristics of each topography. A single St value may characterize the dominant frequency of some flows patterns. However, comparing the full frequency spectra allows for a better understanding of the subtle flow differences among these models. The lift spectra for select models are shown in Figure 14. Each spectra plot is paired with isosurfaces of *Q* criterion, which visualizes the strength of vortices in the wake and demonstrates the variation in these vortical structures.

**Figure 14:**
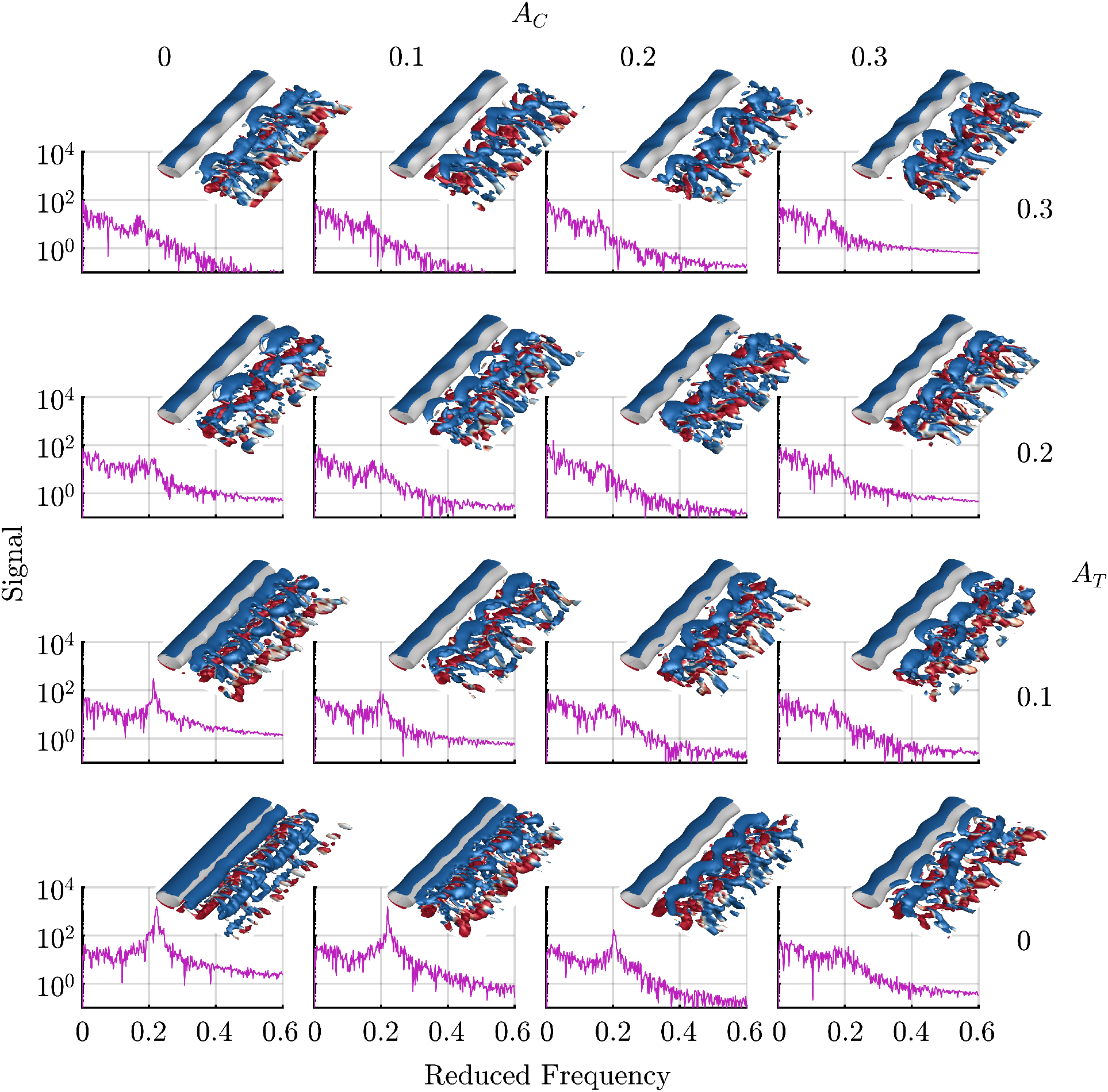
Shedding frequency spectra for a range of cases *A_C_* = 0, 0.1, 0.2, 0.3 and *A_T_* = 0, 0.1, 0.2, 0.3.

The baseline smooth elliptical cylinder, (*A_C_*, *A_T_*) = (0, 0), is shown in the lower left and displays a strong peak in the frequency domain at a reduced frequency of *f*\* = 0.22. This sharp peak is associated with a mostly two-dimensional shedding of alternating sign vorticity typical of smooth cylinders. Moving upwards from the lower left in Figure 14, the dominant amplitude at approximately 0.22 shifts to a slightly lower frequency in (*A_C_*, *A_T_*) = (0, 0.1) before forming lower and broader peaks in (0,0.2) and (0,0.3). This dramatic change is associated with a breakup of the spanwise coherent structures shown in the baseline model and consistent with a large decrease in forces shown in Figure 12. The spectra at (0, 0.3) has a diminished peak at *f*\* = 0.16 with an amplitude approximately two orders of magnitude lower than the dominant peak in the baseline model.

When A_T_ is held constant and AC increases (moving from left to right), the peak gradually reduces in amplitude and slightly decreases its frequency. This is most clearly demonstrated in the bottom two rows. At (*A_C_*, *A_T_*) = (0,0.1) a similar response to the smooth cylinder exists. However it has been replaced with a very broad peak from *f*\* = 0.17 — 0.21 at (0.2,0.1), and only a small peak at *f*\* = 0.17 remains when (*A_C_*, *A_T_*) = (0.3,0.1). The isosurfaces are each unique and show decreasing levels of spanwise coherence with smaller spectra amplitudes and frequency shift. The lowest levels of drag and RMS lift are represented across the top left side of Figure 14, where the amplitude of the signal drops off quickly for any frequencies above the peak at *f*\* = 0.17. At the highest *A_T_* amplitude of 0.3 moving from left to right along the top row, there are fewer changes in the spectra as the dominant frequency is *f*\* = 0.16, shifting to *f*\* = 0.15 when *A_C_* = 0.3. In this row there is a notable increase in amplitude of the higher frequencies with increasing *A_C_*, which is not likely to be visible in the isosurfaces displayed.

## 4. Conclusion

A new surface morphing technique is introduced that is able to modify features of the seal whisker model geometry during CFD simulations, creating a faster and simpler mechanism for exploring geometric variations. Using the base geometric scaffold previously defined by two inclined ellipses [3], the complete surface topography is parameterized, which eliminates small changes in surface rendering between various implementations of seal whisker models. To implement the algorithm, a simple two-dimensional extruded cylinder flow can be used as an initial condition, eliminating the many hours of manual meshing and debugging for complex three-dimensional geometries. Despite the time savings in manual meshing, the mesh morphing algorithm does not increase the overall CPU time.

Next, the mesh morphing algorithm is used to explore the two amplitudes of the seal whisker model, one in the chord direction (*A_C_*) and one in the thickness direction (*A_T_*). Large values of both *A_C_* and *A_T_* (up to 30% of the thickness) are shown to increase spanwise transport although through different mechanisms. The creation of *z*-velocity near the leading edge results in the development of *y*-vorticity, giving large *A_T_* geometries a slight advantage in force reduction compared with large *A_C_* geometries. The combination of extreme values of *A_C_* and *A_T_* does not provide any further reduction in drag or lift forces. At smaller amplitudes, combinations of undulations work together to increase flow three-dimensionality producing local minima such as the model (*A_C_*, *A_T_*) = (0.1, 0.1). Accordingly, the lift force frequency spectra show broader dominant peaks at lower frequencies for cases with considerable breakup of spanwise flow structures.

The results of the 49 model geometries offer much more detail into the force and flow response as a function of undulation amplitude but are consistent with data presented in previous work. The effects of the presence of both undulations were assessed in the computational work by Yoon et al. where, building on the work of Hans et al., seven whisker inspired models were simulated and compared with a smooth elliptical cylinder [13, 17]. Yoon et al. concluded that non-zero values of both *A_C_* and *A_T_* are required for maximum reduction of drag and oscillating lift. They also noted that the introduction of *A_C_* had a larger impact on the force reduction than the introduction of *A_T_* [17]. Within the range of *A_C_* and *A_T_* values simulated, *A_C_* ≈ 0.23 and *A_T_* ≈ 0.09, the results presented here support both of these conclusions. A wider range of amplitude values was investigated by Lui et al., although both amplitude parameters were modified together to maintain a constant local aspect ratio rather than varying *A_C_* and *A_T_* independently [14]. Their results showed consistent low values of 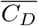 and *C_L,RMS_* across the range of amplitude values simulated. Lyons et al. simulated only two *A_C_* and two *A_T_* values and reported *A_C_* to have a more significant effect on 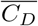 [16]. However in this more detailed exploration, both amplitudes are shown to reduce 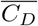 when compared to a smooth elliptical cylinder, albeit through different mechanisms.

This newly proposed mesh transformation algorithm can further enable feature variation within the seal whisker inspired geometries as they apply to various engineering applications in drag reduction, vibration suppression, or hydrodynamic sensing. Although not explored here, future work could include active flow control techniques enabled through the mesh morphing algorithm.

## 5. Acknowledgement

The authors acknowledge Brown University’s Center for Computation and Visualization for computational resources. Financial support is graciously provided by University of Wisconsin-Madison’s Hilldale Undergraduate/Faculty Research Fellowship, Wisconsin Alumni Research Foundation’s Fall Research Competition, and the National Science Foundation Award 2035789. Thank you to Kirby Heck and Trevor Dunt for suggestions to the manuscript.

## Appendix A. Appendix

### Appendix A.1. Derivation of R_1_ and R_2_

An ellipse with radii *a* and *b*, can be expressed with the equation

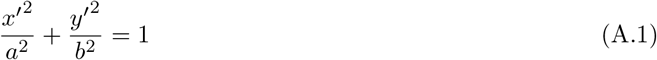

in the coordinate system *x*′, *y*′ that has an angle α with respect to the *x*-axis.

In order to complete calculations parallel and perpendicular to the fluid flow, the coordinate system can be rotated about the *y*-axis using the transform

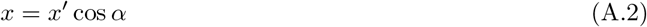

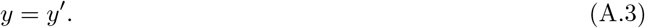

The ellipse is now represented in terms of *x* and *y* as

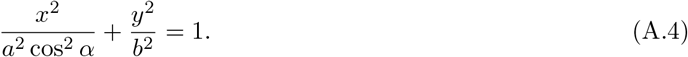

Next, substituting *y* = *x* tan *θ*, and solving for *x*, the ellipse equation can be rewritten in terms of *θ* as

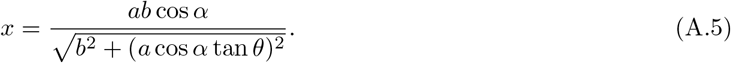

Then, *R*_1_ is defined by

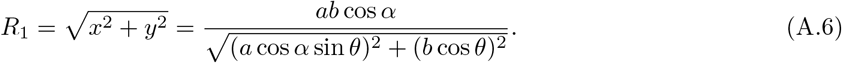

Using the same procedure, *R*_2_ can be also derived as

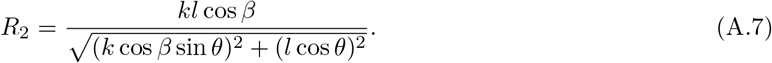

